# GPR35 inhibits TRPA1-mediated colonic afferent hypersensitivity through suppression of Substance P release

**DOI:** 10.1101/2023.09.12.557415

**Authors:** Rohit A Gupta, James P Higham, Paulina Urriola-Muñoz, Katie H Barker, Luke Paine, Joshua Ghooraroo, James RF Hockley, Taufiq Rahman, Ewan St John Smith, Alistair JH Brown, Rie Suzuki, David C Bulmer

**Author notes:** Corresponding author: David C Bulmer.

## Abstract

The development of non-opioid analgesics for the treatment of chronic abdominal pain is a pressing area of unmet clinical need. To address this, we examined the expression of G_i/o_-coupled receptors in colonic sensory neurons, which, like opioid receptors, have the potential to inhibit nociceptor activation due to their inhibitory G protein coupling. This led to the identification of the orphan receptor GPR35 as a visceral analgesic drug target due to its marked co-expression with TRPA1, a mediator of noxious mechanotransduction in the bowel. Consistent with *in silico* docking studies which identified binding sites for the mast cell stabiliser cromolyn and phosphodiesterase inhibitor zaprinast at GPR35, we demonstrated, using GPR35 knockout mice, that the antinociceptive effects of these drugs on TRPA1-mediated colonic nociceptor activation and mechanosensitisation were GPR35-dependent. Further work showed these antinociceptive effects occurred through the inhibition of substance P (SP) release. This confirmed both the pronociceptive effect of SP on colonic afferents, and the contribution of SP to TRPA1-mediated colonic nociceptor activation and sensitisation. We also found that TRPA1-induced contraction of the colon was mediated by SP signalling and could be inhibited by cromolyn in a GPR35-dependent manner. Our data identify GPR35, through its inhibition of SP-mediated colonic contractility and nociceptor activation and sensitisation, as a putative mechanism for the reported clinical efficacy of cromolyn in the treatment of irritable bowel syndrome. These findings highlight the potential utility of GPR35 agonists to deliver non-opioid analgesia for the treatment of abdominal pain associated with gastrointestinal diseases such as irritable bowel syndrome and inflammatory bowel disease.

## Introduction

The development of non-opioid analgesics to treat chronic pain is a clinical priority due to the risk of dependence and addiction associated with long-term opioid use (Woodcock, 2009; Barnes, 2020). This is particularly pressing for the treatment of abdominal pain evoked by gastrointestinal diseases in which the use of both opioid (Burr et al., 2018; and Cohen-Mekelburg et al., 2018) and non-steroidal anti-inflammatory drugs (Wolfe et al., 1999; Takeuchi et al., 2006) is further contraindicated due to gut-specific side effects, such as constipation and gastrointestinal bleeding, respectively. Given the high prevalence of nociplastic abdominal pain evoked by conditions such as irritable bowel syndrome (IBS) and the rising incidence of chronic pain due to inflammatory bowel disease (IBD), there is a considerable unmet clinical need for development of viscerospecific analgesics to address the marked personal and socioeconomic burden associated with chronic abdominal pain (Zeitz et al., 2016; Perler et al., 2019; Ford et al., 2017).

The development of visceral hypersensitivity in response to mediators released from the bowel of patients with IBS or IBD is a key pathophysiological event, in which visceral nociceptors become sensitised and therefore signal pain in response to previously innocuous distension or contraction of the bowel (Bueno and Fioramonti, 2002; Gold and Gebhart, 2010). Numerous mechanisms contribute to the development of visceral hypersensitivity, such as enhanced activity of ion channels which mediate mechanosensation in the bowel, such as transient receptor potential ankyrin 1 (TRPA1) channels (Brierley et al., 2009). Moreover, mechanosensitive afferent excitability can also be increased due to depolarisation of their resting membrane potential or reduced threshold for action potential firing following modulation of ion channels, such as the voltage gated sodium channels Na_V_1.8 (Huang et al., 2013) and Na_V_1.9 (Hockley et al., 2014). The mediators responsible for visceral hypersensitivity are not fully understand, although mast cell derived mediators such as histamine, serotonin (5-HT), nerve growth factor and tryptase, as well as lipids such as 5-oxo-ETE, are implicated in IBS, in addition to immune mediators such as cytokines (e.g., tumour necrosis factor alpha, TNFα) and proteinases (e.g., neutrophil elastase) being implicated in IBD (Barbara et al., 2007; Cenac et al., 2007; Balemans et al., 2019 and Barker et al., 2022). These mediators and mechanisms therefore represent novel drug targets for the development of visceral analgesics, although there is a risk of redundancy when targeting a single sensitising agent.

An alternative approach is to stimulate G_i/o_-coupled receptors, which have the potential to inhibit nociceptor activation irrespective of the noxious stimulus due to their downstream effector responses that can enhance potassium channel conductance, reduce the pro-nociceptive effects associated with cAMP accumulation and suppress neuropeptide release (Yudin and Rohacs, 2018; Kano et al., 2019). G_i/o_-coupled receptors comprise a broad range of members stimulated by agonists with analgesic activity, such as opioids and cannabinoids. To identify putative analgesic approaches for the treatment of abdominal pain in gastrointestinal disease, we compared the co-expression of transcripts for G_i/o_-coupled receptors with TRPA1, a mediator of colonic nociceptor mechanosensation, using our existing single cell RNA-sequencing colon sensory neuron transcriptomic data (Hockley et al., 2019). This led to the identification of the orphan G-protein-coupled receptor, GPR35, as a novel non-opioid visceral analgesic drug target. Consequently, we evaluated the effect of GPR35 agonists on TRPA1-mediated colonic nociceptor activation, mechanical hypersensitivity, colonic contractility and neuropeptide release. This work demonstrates an anti-nociceptive role for GPR35 agonists, such as cromolyn, and thereby highlights potential as visceral analgesics.

## Methods

### Ethical approval

Experiments using animals or animal tissue were conducted in compliance with the Animals (Scientific Procedures) Act 1986 Amendment Regulations 2012 under Project Licences P7EBFC1B1 and PP5814995 granted to E. St. J. Smith by the Home Office with approval from the University of Cambridge Animal Welfare Ethical Review Body. Animals were euthanised by rising concentration of CO_2_ or cervical dislocation followed by exsanguination.

### Molecular modelling and docking

A homology model of mouse GPR35 receptor (mGPR35, Uniprot id: Q9ES90) was built using the published structure of human GPR35 receptor (hGPR35R, PDB id: 8H8J) using SwissModel server (https://swissmodel.expasy.org/). The initial model of mGPR35 was then further refined through GalaxyRefine 2 module of the GalaxyWEB server (https://galaxy.seoklab.org/). The refined mGPR35 model was used for docking alongside the published structure of hGPR35.

Docking of the chosen ligands were carried out against the unliganded hGPR35 and modelled mGPR35R structures using published protocol (Callejo et al., 2020). Briefly, the 3D structures of zaprinast, cromylyn and lodoxamide were obtained from PubChem and first blindly docked to the GPR35 structures using AutoDock 4.2.6. The best poses of the two drugs were then used for a focused docking using GOLD suite version 5.3 (CCDC, Cambridge) using 10 independent docking runs. The 2D ligand interaction diagrams of the final docked poses were generated through PoseView™ implemented in the ProteinsPlus webserver (https://proteins.plus/). All relevant images were produced in PyMol version 1.3.

### Animals

Adult male C57BL/6 mice (8-16 weeks) or CD1 mice (8-16 weeks) were obtained from Charles River (Cambridge, UK; RRID:IMSR_JAX:000664) and male GPR35^-/-^ mice (C57BL/6-Gpr35tm1b(EUCOMM)Hmgu/WtsiH) rederived by MRC Harwell. Mice were housed in temperature-controlled rooms (21°C) with a 12-hour light/dark cycle and provided with nesting material, enrichment (e.g., tubes, chewing blocks and shelters) and access to food and water *ad libitum*.

### *Ex-vivo* whole nerve electrophysiology

#### Electrophysiological recording

Electrophysiological recordings of lumbar splanchnic nerve (LSN) activity were conducted as previously described (Peiris et al., 2017; Barker et al., 2022). Briefly, the distal colorectum (splenic flexure to rectum) and associated LSN (rostral to inferior mesenteric ganglia) were isolated from mice euthanised as described above. The colon was cannulated with fine thread (Gutermann) in a rectangular Sylgard-coated recording chamber (Dow Corning, UK) and bath superfused (7 ml/min; 32-34°C) and luminally perfused (100 μL/min) by a syringe pump (Harvard apparatus, MA) against a 2-3mmHg with carbogenated Krebs buffer solution (95% O_2_, 5% CO_2_). Krebs buffer was supplemented with 10 μM atropine and 10 μM nifedipine to prevent smooth muscle activity.

Borosilicate suction electrodes were used to record the multi-unit activity of LSN bundles. Signals were amplified (gain 5 kHz), band pass filtered (100–1300 Hz; Neurolog, Digitimer Ltd, UK), and digitally filtered for 50 Hz noise (Humbug, Quest Scientific, Canada). Analogue signals were digitized at 20 kHz (Micro1401; Cambridge Electronic Design, UK) and signals were visualised with Spike2 (Cambridge Electronic Design, UK). Preparations underwent a minimum 30-minute stabilisation period to ensure baseline firing was consistent.

Pilot studies examining the effect of the TRPA1 agonist ASP7663 on colonic afferent activity used tissue from CD1 mice. These studies examined the effect of a repeated 20 ml bath perfusion with 100 μM ASP7663 (final bath concentration 30 μM) at 45-minute intervals confirming the marked tachyphylaxis to repeat application. Furthermore, separate experiments examined the concentration dependency of the ASP7663 response by 20 ml bath perfusion with ASP7663 at 30 μM, or 10 μM (final bath concentration of 9 μM and 3 μM respectively). The contribution of TRPA1 to ASP7663 responses was examined by bath perfusion with 20 ml ASP7663 30 μM (bath concentration 9 μM) in the presence of the TRPA1 antagonist AM0902 (1 μM), given immediately following pre-treatment by bath perfusion with 100 ml AM0902 (1 μM).

Further studies were performed using tissue from wildtype C57BL/6 mice and GPR35^-/-^ mice. For TRPA1 agonist mediated colonic afferent activation protocols, the effect of pre-treatment with either 100 ml vehicle (0.1% DMSO), 100ml cromolyn (1 μM, 10 μM or 100 μM) or 50 ml zaprinast (10 μM or 100 μM) were examined on the colonic afferent response to bath perfusion with 20ml ASP7663 100 μM (bath concentration 30 μM) given in the presence of respective vehicle, cromolyn or zaprinast concentrations in tissue from wildtype mice. While the effects of pre-treatment with either 100 ml vehicle (0.1% DMSO), 100 ml cromolyn (100 μM) or 50 ml zaprinast (100 μM) were also examined on the colonic afferent response to bath perfusion with 20 ml ASP7663 100 μM (bath concentration 30 μM) given in the presence of respective vehicle, cromolyn or zaprinast concentrations in tissue from GPR35^-/-^ mice.

In studies examining the effect of mast cell degranulation on colonic afferent firing, the response to bath perfusion with 100 ml of compound 48/80 (50 µg/ml) was examined in tissue from wildtype C57BL/6 mice. Studies examining the effect of phosphodiesterase (PDE) inhibition on the colonic afferent response to ASP7663 measured the effect of pre-treatment with either 100 ml of the pan PDE inhibitor IBMX (50 μM) or 100 ml of the PDE5/6 selective inhibitor sildenafil (1 μM) on the colonic afferent response to 20 ml ASP7663 100 μM (bath concentration 30 μM) given in the presence of respective PDE inhibitor.

For studies examining the afferent response to luminal distension and its sensitisation by ASP7663. Repeated ramp distensions (0-80 mmHg) were performed by occluding the luminal perfusion outflow which gradually increased intraluminal pressure up to 80 mmHg over approximately 1 to 2 minutes. When the maximum 80 mmHg pressure was achieved, the luminal outflow was re-opened, allowing pressure return to baseline. Pilot studies examined the effect of repeat luminal distension applied at 10 min intervals, confirming that although a modest reduction in the colonic afferent response to distension can be observed between the first and second repeat distension the afferent response to the second and third repeat distension was comparable. As such the effect of pre-treatment with vehicle (0.1 DMSO, 100 ml), AM0902 (3 μM, 100 ml), cromolyn (100 μM, 100 ml) and zaprinast (100 μM, 50 ml) was examined on the afferent response to the third distension. Pre-treatment with zaprinast began 5-minutes before the third distension (total interval time 10-minutes), all other treatments began 10-minutes before the third distension. For studies looking at the effect of zaprinast and cromolyn on the sensitisation of colonic afferent responses to luminal distension by ASP7663, pre-treatment with either 50 ml zaprinast (10 and 100 μM), 100 ml cromolyn (1, 10 and 100 μM) or respective 50 ml or 100 ml vehicle (0.1% DMSO) began immediately following the second ramp distension, after which the bath was perfused with 20 ml of ASP7663 100 μM (bath concentration 30 μM) in the presence of respective pre-treatments and the third luminal distension applied 10-minutes later. This experimental protocol was repeated in tissue from GPR35^-/-^ mice for the 100 ml vehicle (0.1% DMSO) and 100 ml cromolyn (100 μM) pre-treatment.

For studies looking at the contribution of substance P (SP) to colonic afferent firing and response to ASP7663, we first examined in separate experiments, the effect of 20 ml perfusion with ASP7663 100 μM (bath concentration 30 μM) alone, and in the presence of the neurokinin 1 receptor (NK_1_R) antagonist aprepitant (10 μM) following a 100 ml pre-treatment with aprepitant (10 μM). Following which we also studied in separate experiments the effect of bath perfusion with 20 ml SP at 10 μM (bath concentration 3 μM), 30 μM (bath concentration 9 μM), 50 μM (bath concentration 15 μM) concentration. In addition to examining the effect of bath perfusion with 20 ml SP at 50 μM (bath concentration 15 μM) in the presence of thiorphan (10 µM) and captopril (10 µM), following a 100 ml pre-treatment with thiorphan (10 µM) and captopril (10 µM) alone and repeated in the presence of aprepitant (10 µM).

Finally, we examined the contribution of SP to the afferent response to distension (0-80 mmHg) and its sensitisation by ASP7663. To do this we examined the colonic afferent response to a third repeat luminal distension given 10-minutes after a 120 ml pre-treatment with with thiorphan (10 µM) and captopril (10 µM) alone started immediately following the second repeat luminal distension (given 10-minutes after the first), or 10-minutes following a 100 ml pre-treatment with thiorphan (10 µM) and captopril (10 µM), and a 20 ml perfusion with SP 50 μM (bath concentration 15 μM) in the presence of thiorphan (10 µM) and captopril (10 µM). In addition, we also examined the colonic afferent response to a third luminal distension applied 10-minutes after a 100 ml pre-treatment with either aprepitant (10 μM) or vehicle (0.1% DMSO) and subsequent 20 ml application of ASP7663 100 μM (bath concentration 30 μM) given in the presence of respective pre-treatments. Finally, the difference in the colonic afferent response to luminal distension was compared between a second repeat luminal distension and a third repeat distension given following a 100 ml perfusion with aprepitant (10 μM).

#### Analysis

LSN discharge was determined by quantifying the number of spikes passing a manually-determined threshold typically set at twice the background noise (60-80 µV) and binned to determine the firing rate (spikes/s). The change in firing rate was found by subtracting baseline activity (average firing rate in the 3-minutes preceding the stimulus) from increases in nerve activity following stimulus application. To quantify mechanical hypersensitivity following drug application, the change in afferent firing during ramp 3 (post-drug) was compared with the changes in afferent firing during ramp 2 (pre-drug) for individual treatments. To compare the change in mechanical hypersensitivity between vehicle or treatments, post-drug responses (ramp 3) were normalised to pre-drug responses. As such, a peak percentage change in afferent firing during ramp 3 of ∼100% indicates no change in distension-evoked firing following treatment and hence, no induction of mechanical hypersensitivity.

### Ca^2+^ imaging

#### Primary culture of dorsal root ganglia neurons

Cells from the dorsal root ganglia (DRG) were cultured as described previously (Barker et al., 2022). In brief, the spine was removed and bifurcated to allow isolation of DRG. DRG innervating the distal colon (T12-L5) were removed into ice-cold Leibovitz’s L-15 GlutaMAX™ medium (supplemented with 2.6% (v/v) NaHCO_3_). DRG were then incubated with 1 mg/mL collagenase (15-minutes) followed by 1 mg/ml trypsin (30-minutes), both with 6 mg/ml bovine serum albumin (BSA) in Leibovitz’s L-15 GlutaMAX™ medium (supplemented with 2.6% (v/v) NaHCO_3_). Following removal of the enzymes, DRG were resuspended in 2 ml Leibovitz’s L-15 GlutaMAX™ medium containing 10% (v/v) foetal bovine serum (FBS), 2.6% (v/v) NaHCO_3_, 1.5% (v/v) glucose and 300 units/ml penicillin and 0.3 mg/ml streptomycin (P/S). DRG were mechanically triturated using a 1 ml Gilson pipette and centrifuged (100 *g*) for 30-seconds. The supernatant, containing dissociated DRG neurons, was collected in a separate tube and the pellet resuspended in 2 ml Leibovitz’s L-15 GlutaMAX™ medium containing 10% (v/v) FBS, 2.6% (v/v) NaHCO_3_, 1.5% (v/v) glucose and P/S. This process was repeated five times, after which the total supernatant was centrifuged (100 *g*) for 5-minutes to pellet the dissociated DRG neurons. Cells were resuspended in 250 µl Leibovitz’s L-15 GlutaMAX™ medium containing 10% (v/v) FBS, 2.6% (v/v) NaHCO_3_, 1.5% (v/v) glucose and P/S and plated (50 µl per dish) onto 35 mm glass bottomed dishes coated with poly-D-lysine (MatTek, MA, USA) and laminin (Thermo Fisher: 23017015). Dishes were incubated for 3-hours (37°C, 5% CO_2_) to allow cell adhesion and subsequently flooded with Leibovitz’s L-15 GlutaMAX™ medium containing 10% (v/v) FBS, 2.6% (v/v) NaHCO_3_, 1.5% (v/v) glucose and P/S and incubated overnight. Neurons were used for experiments after 16-24 hours in culture.

#### Imaging

Extracellular bath solution (in mM: 140 NaCl, 4 KCl, 1 MgCl_2_, 2 CaCl_2_, 4 glucose, 10 HEPES) was prepared and adjusted to pH 7.4 using NaOH and an osmolality of 290-310 mOsm with sucrose. Culture medium was aspirated from neuronal cultures and cells were incubated (30-minutes) with 100 µl Fluo-4-AM (10 μM) diluted in bath solution (room temperature; shielded from light). Prior to imaging, Fluo-4-AM was removed, and dishes were washed with bath solution.

Dishes were mounted on the stage of an inverted microscope (Nikon Eclipse TE-2000S) and cells were visualised with brightfield illumination at 10x magnification. To ensure a rapid exchange of solutions during protocols, the tip of a flexible perfusion inflow tube (AutoMate Scientific, CA, USA) attached to a valve-controlled, gravity-fed perfusion system (Warner Instruments, CT, USA) was placed beside the field of view. Cells were superfused with bath solution to establish a baseline.

Fluorescent images were captured with a charge-coupled device camera (Retiga Electro, Photometrics, AZ, USA) at 2.5 fps with 100 ms exposure. Fluo-4 was excited by a 470 nm light source (Cairn Research, Faversham, UK). Emission at 520 nm was recorded with μManager. All protocols began with a 10-second baseline of bath solution before drug superfusion (SP 100 nM) followed by capsaicin (1 µM) after a 4-minute wash period with bath solution. Finally, cells were stimulated with 50 mM KCl for 10-seconds to determine cell viability, identify neurons and allow normalisation of fluorescence.

#### Analysis

Image analysis was carried out using ImageJ (NIH, MA, USA). Regions of interest were traced around individual cells. Average pixel intensity across each region of interest was measured and analysed with custom-written scripts in RStudio (RStudio, MA, USA) to compute the proportion of neurons stimulated by each drug application. Briefly, the background fluorescence was subtracted from each region of interest, and fluorescence intensity (F) was baseline-corrected and normalised to the maximum fluorescence elicited during the application of 50 mM KCl (F_pos_). Maximal fluorescence in KCl was denoted as 1 F/F_pos_. Further analysis was confined to cells with a fluorescence increase ≥5% above the mean baseline before 50 mM KCl application, as these were considered neurons. Using manual quality control, neurons were deemed responsive to drug application if a fluorescence increase of 0.1 F/F_pos_ was observed following drug superfusion. Responses to drug application were discounted if the increase in fluorescence began before or after the period of drug superfusion. Recordings without a stable baseline were also excluded from analysis. The proportion of drug-responsive neurons (i.e., proportion of KCl-sensitive neurons exhibiting >0.1 F/F_pos_ increase in fluorescence during drug application) was measured for each experiment.

### Chemiluminescent immunoassay to measure substance P (SP) release

The distal colon from wildtype and GPR35^-/-^ mice was removed, cleared of faecal material, and bathed with oxygenated Krebs solution. The tissue was washed 3 times with Krebs buffer before the experiment to remove any adhering tissue debris or blood that may interfere with the experiment. A segment of the colon (∼2 cm in length) was transferred into filtered carbogenated Krebs buffer and equilibrated at 37°C for 15-minutes. Tissue segments were then incubated with ASP7663 (100 µM) alone or following pre-incubation with either cromolyn (100 µM) or AM0902 (3 µM) for 15-minutes. All incubations were carried out in the presence of thiorphan and captopril to prevent the breakdown of SP. Following this, the solution bathing the colon was removed and centrifuged (500 *g* for 5-minutes), with the resulting supernatant used to measure SP with a commercially-available competitive chemiluminescent immunoassay (CLIA) kit according to the manufacturer’s instructions (Cloud-Clone Corp., TX, USA, Cat. no. CCA393Mu). The SP concentration is each sample was found through the construction of standard curves, and SP concentration was normalised to tissue weight.

### Colonic contractility assay

The distal colon (∼2 cm) was removed from wildtype and GPR35^-/-^ mice, cleared of faecal material and bath in ice-cold carbogenated Krebs buffer until use. The colon was mounted vertically in a tissue bath filled with carbogenated Krebs buffer maintained at 37°C, with one end of the colon attached to an isotonic transducer which converted changes in colon length into voltages. Signals were displayed using LabChart (ADInstruments). Tissue equilibrated for 60-minutes (washed every 15-minutes) prior to starting experimental protocols.

To assess baseline tissue contractility, 100 µM acetylcholine was applied. After washout, ASP7663 (100 µM) was applied alone or following a 10-minute pre-incubation with either AM0902 (3 µM), aprepitant (10 µM) or cromolyn (100 µM). At the end of each protocol, tissues were washed and 100 µM acetylcholine was re-applied to ensure tissue contractility was unaffected and the tissue had not been paralysed. The response to ASP7663 in each condition was normalised to the peak response evoked by the first application of acetylcholine.

### Statistics

All datasets were scrutinized to ensure they met the assumptions (e.g., normality assessed using Shapiro-Wilk test) of parametric analyses and, where appropriate, rank-based non-parametric analyses were employed. Detailed of statistical tests used in each experiment are in the relevant figure legends. Data are displayed as mean ± standard error. P-value cutoffs in figures are denoted by *p<0.05, **p<0.01 and ***p<0.001.

## Results

### GPR35 is highly co-expressed with TRPA1 in colonic sensory neurons

To identify novel visceral analgesics, we examined the co-expression of G_i/o_-coupled receptors with TRPA1, an effector of noxious colonic mechanosensation, in colonic neurons using our previously published transcriptomic profiling of colon-innervating sensory neurons (Figure 1A). Findings from this analysis identified a cluster of G_i/o_-coupled receptors highly co-expressed with TRPA1 in colonic neurons. These included the orphan receptor GPR35 (95.1% of *Trpa1*-positive neurons also expressed *Gpr35)* and receptors with known (or experimentally demonstrated) analgesic properties, such as the µ opioid receptor (*Oprm1*, 93.2%), the type 1 cannabinoid receptor (*Cnr1*, 95.1%), the adenosine A1 receptor (*Adora1*, 98.5%) and the GABA_B1_ receptor (*Gabbr1*, 100%). GPR35 activation is not linked to central side effects, such as sedation, associated with opioid (Del et al., 2017), cannabinoid (Huestis et al., 2019), or GABA receptor agonists (McKernan et al., 2000), or to cardiovascular side effects associated with adenosine receptor agonists (Mustafa et al., 2009). Both *Trpa1* and *Gpr35* transcripts were found to be highly expressed in colonic sensory neurons (Figure 1B), with *Gpr35* broadly expressed across all colonic afferent populations, while *Trpa1* was more selectively expressed in peptidergic afferent populations (Figure 1C). Given the high co-expression of *Gpr35* and *Trpa1* (Figure 1D), we chose to further investigate the role of this receptor in the regulation of colonic mechanotransduction.

**Figure 1:**
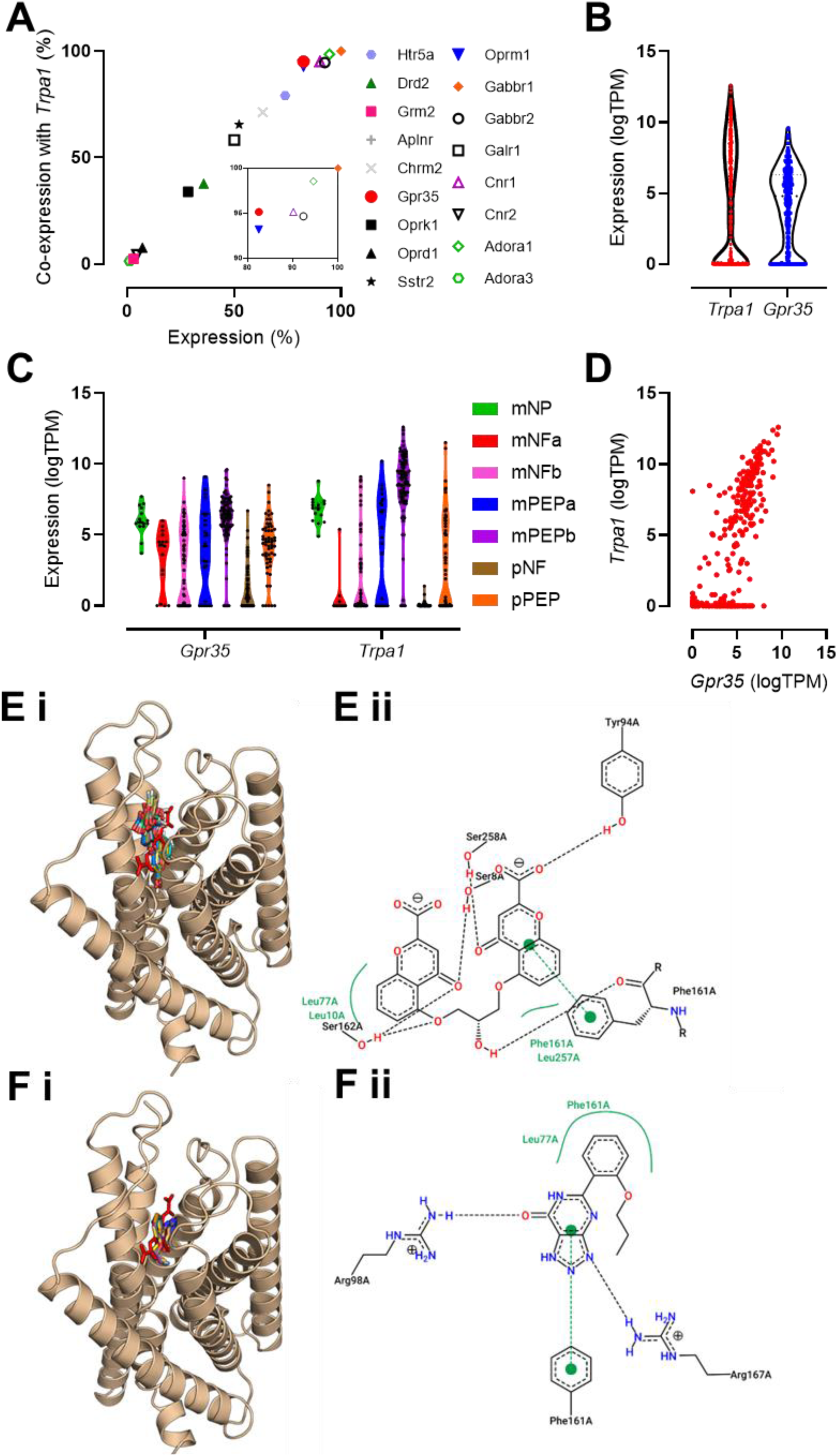
Expression of GPR35 in colonic sensory neurons and validation of cromolyn and zaprinast as GPR35 ligands. (A) Co-expression of transcripts encoding selected G_i/o_-coupled receptors with *Trpa1* in colonic sensory neurons. *Inset*: enlargement of top-right cluster of data points. Data in – (D) redrawn from Hockley *et al*., 2019. (B) Expression (log[transcripts per million]) of transcripts encoding *Trpa1* and *Gpr35* in colonic sensory neurons. (C) Expression of transcripts encoding *Trpa1* and *Gpr35* in each subpopulation of colonic sensory neuron. NP, non-peptidergic; NF, neurofilament-expressing; PEP, peptidergic; m, mixed lumbar splanchnic and pelvic afferents; p, pelvic afferents. (D) Co-expression of transcripts encoding *Trpa1* and *Gpr35* in colonic sensory neurons. (E) *(i)* Predicted binding mode of cromolyn at mouse GPR35 based on the ten best ranked binding poses obtained through independent docking runs in GOLD v.5.3. *(ii)* Ligand interaction diagram generated using PoseView. The most representative pose for cromolyn is shown. Dotted black lines indicate hydrogen bonding, solid green lines represent hydrophobic interactions and dotted green lines represent π-π stacking. (F) *(i)* Predicted binding mode of zaprinast at mouse GPR35 based on the ten best ranked binding poses obtained through independent docking runs in GOLD v.5.3. *(ii)* Ligand interaction diagram generated using PoseView. The most representative pose for zaprinast is shown. Dotted black lines indicate hydrogen bonding, solid green lines represent hydrophobic interactions and dotted green lines represent π-π stacking.

**Supplemental Figure 1:**
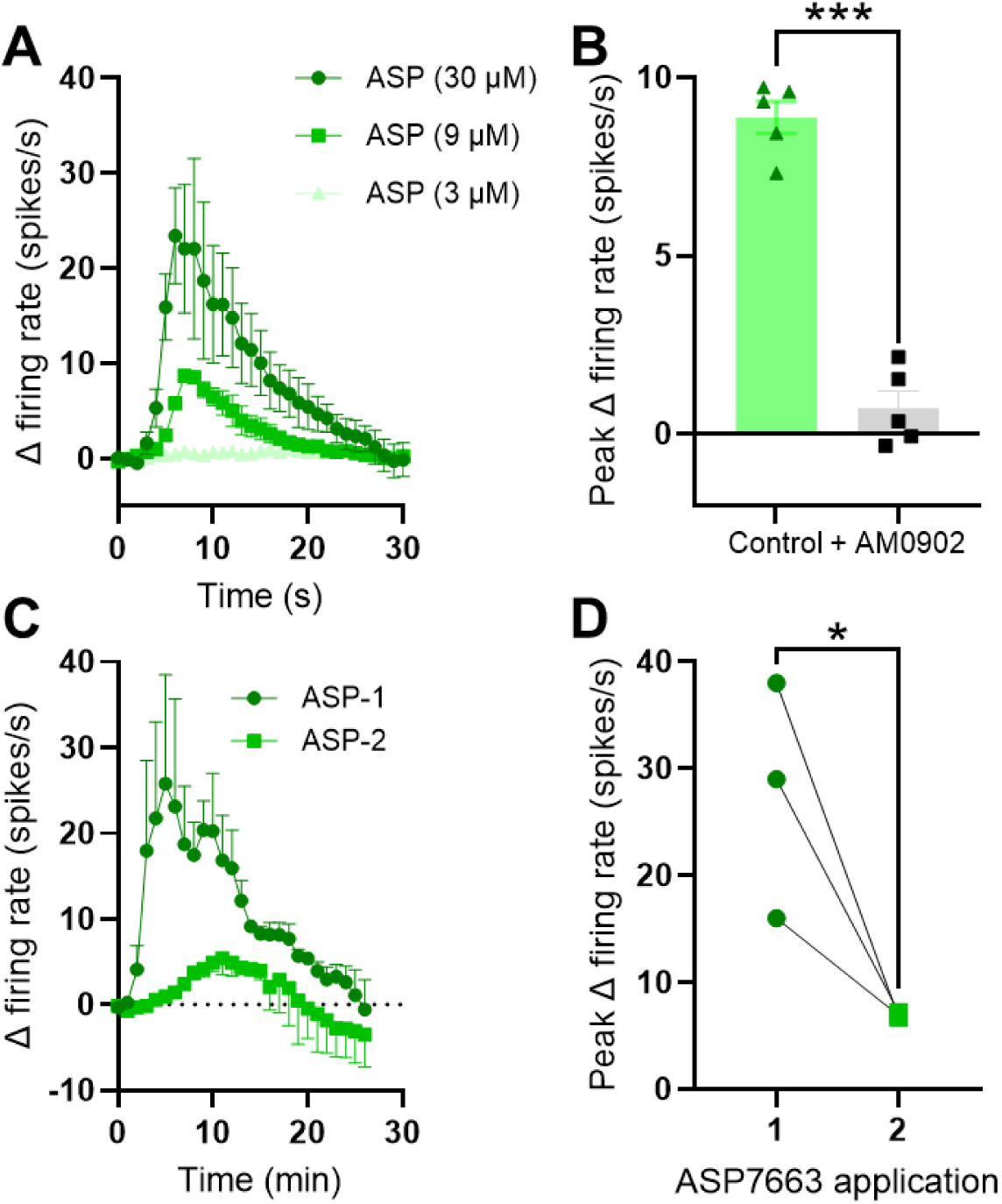
ASP7663 stimulates colonic afferent firing through TRPA1 activation. (A) Grouped data showing the mean change in afferent firing following the application 3, 9 or 30 µM ASP7663. Symbols show the mean firing rate and error bars show the standard error. (B) Grouped data showing the mean peak change in afferent firing rate following the application of 9 µM ASP7663 to tissue in control conditions or pre-incubated with the TRPA1 antagonist AM0902 (1 µM). Two-tailed unpaired t-test. (C) Grouped data showing the mean change in afferent firing following the first and second application of ASP7663. (D) Grouped data showing the mean peak change in afferent firing following the first and second application of ASP7663. Two-tailed ratio-paired t-test.

We performed *in silico* semi-rigid blind docking followed by ligand pose refinement to predict the binding mode of two GPR35 agonists, cromolyn (Yang et al., 2010; Jenkins et al., 2010 and 2012) and zaprinast (Taniguchi et al., 2006; Guo et al., 2008; Jenkins et al., 2010 and 2012), at mouse GPR35 to determine their suitability for use in functional studies. Both cromolyn (Figure 1Ei) and zaprinast (Figure 1Fi) appear to bind at the orthosteric site on mouse GPR35, where lodoxamide, a synthetic GPR35 agonist, is known to bind (Duan et al., 2022). There are clear differences in the predicted residues involved in the binding of cromolyn and zaprinast to mouse GPR35, e.g. X and Y, although π-π with Phe161 appears to be important for the interaction of both ligands with the receptor (Figures 1Eii and 1Fii).

### TRPA1-mediated colonic afferent activation is inhibited by stimulation of GPR35

Following the identification of GPR35 as a putative visceral analgesic drug target, we sought to determine the ability of GPR35 agonists to prevent the activation of colonic afferents by the noxious TRPA1-selective agonist, ASP7663 (Ko et al., 2019).

Pilot studies carried out in tissue from CD1 mice confirmed that ASP7663 elicited a TRPA1-mediated increase in colonic afferent fibre activity. These data showed that ASP7663 evoked a concentration dependent increase in afferent firing of 0.6±0.1 spikes/s at 3 µM (N = 5), 8.9±0.5 spikes/s at 9 µM (N = 5) and 29.4±7.0 spikes/s at 30 µM (N = 5), respectively (Supplemental Figure 1A). The afferent response to 9 µM ASP7663 was abolished by treatment with the TRPA1-selective antagonist, AM0902 (1 µM) (N = 5, p < 0.0001, Supplemental Figure 1B). Afferent responses to ASP7663 underwent marked tachyphylaxis on repeat application of ASP7663 at 30 µM (Supplemental Figure 1C and D), limiting studies with GPR35 agonists to a single application of ASP7663.

To test whether activation of GPR35 could inhibit the response to ASP7663 (Figure 2A), tissue from C57BL/6 mice was pre-incubated with increasing concentrations of cromolyn sodium (CS, Figure 2B) or zaprinast (Zap, Figure 2C). Pre-treatment with CS (1 µM) failed to attenuate ASP7663 (30 µM)-induced afferent firing (13.1±2.8 spikes/s [N = 6] vs 9.9±1.7 spikes/s [N = 5], p = 0.64). However, pre-incubation with 10 µM CS did suppress afferent activity following ASP7663 application to 4.1±1.3 spikes/s (p = 0.0065). 100 µM CS inhibited the afferent response to ASP7663 to a similar extent (4.8±1.8 spikes/s, N = 5, p = 0.02, Figure 2D and E). Pre-incubation with 10 µM Zap did not affect ASP7663-induced afferent firing (12.0±1.6 spikes/s, N = 5, p = 0.99), though treatment with 100 µM Zap substantially reduced the effect of ASP7663 on afferent firing (3.4±1.5 spikes/s, N = 5, p = 0.0051, Figure 2D and E).

**Figure 2:**
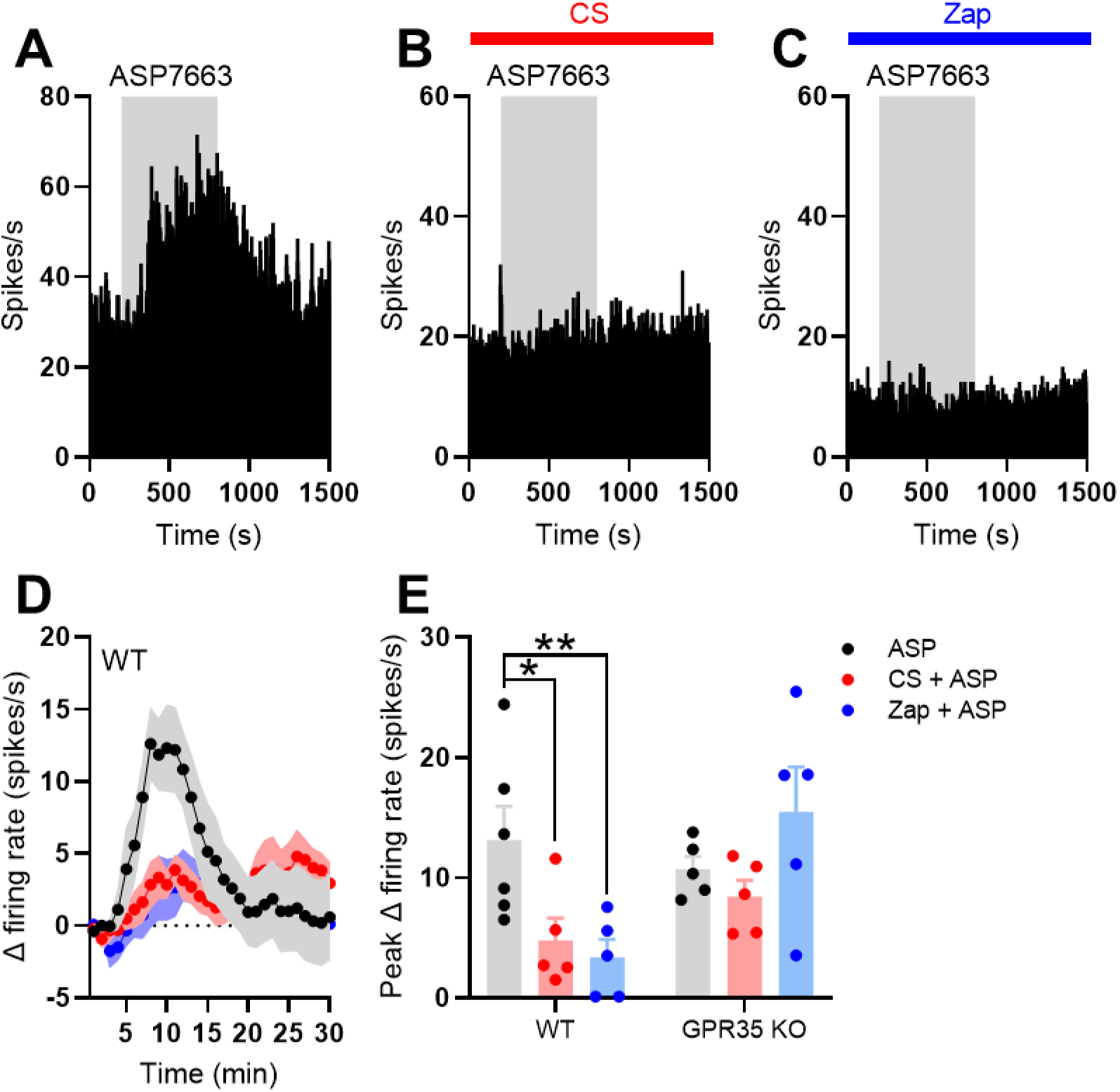
TRPA1-mediated colonic afferent firing is attenuated by activation of GPR35. (A) Example rate histogram showing the change in afferent firing evoked by the application of ASP7663 (grey shaded region). (B)Example rate histogram showing the change in afferent firing evoked by the application of ASP7663 (grey shaded region) in the presence of cromolyn (CS, 100 µM, red bar). (C) Example rate histogram showing the change in afferent firing evoked by the application of ASP7663 (grey shaded region) in the presence of zaprinast (Zap, 100 µM, blue bar). (D) Grouped data showing the mean change in afferent firing evoked by ASP7336 (points indicate the mean; shaded regions show the standard error) in the presence of either CS (100 µM, red) or Zap (100 µM, blue). (E) Grouped data showing the peak change in afferent firing evoked by ASP7336 in the presence of either CS (100 µM, red) or Zap (100 µM, blue) in both wildtype (WT) and GPR35^-/-^ animals. Data for each genotype analysed using a one-way ANOVA with Dunnett’s post-hoc tests.

**Supplemental Figure 2:**
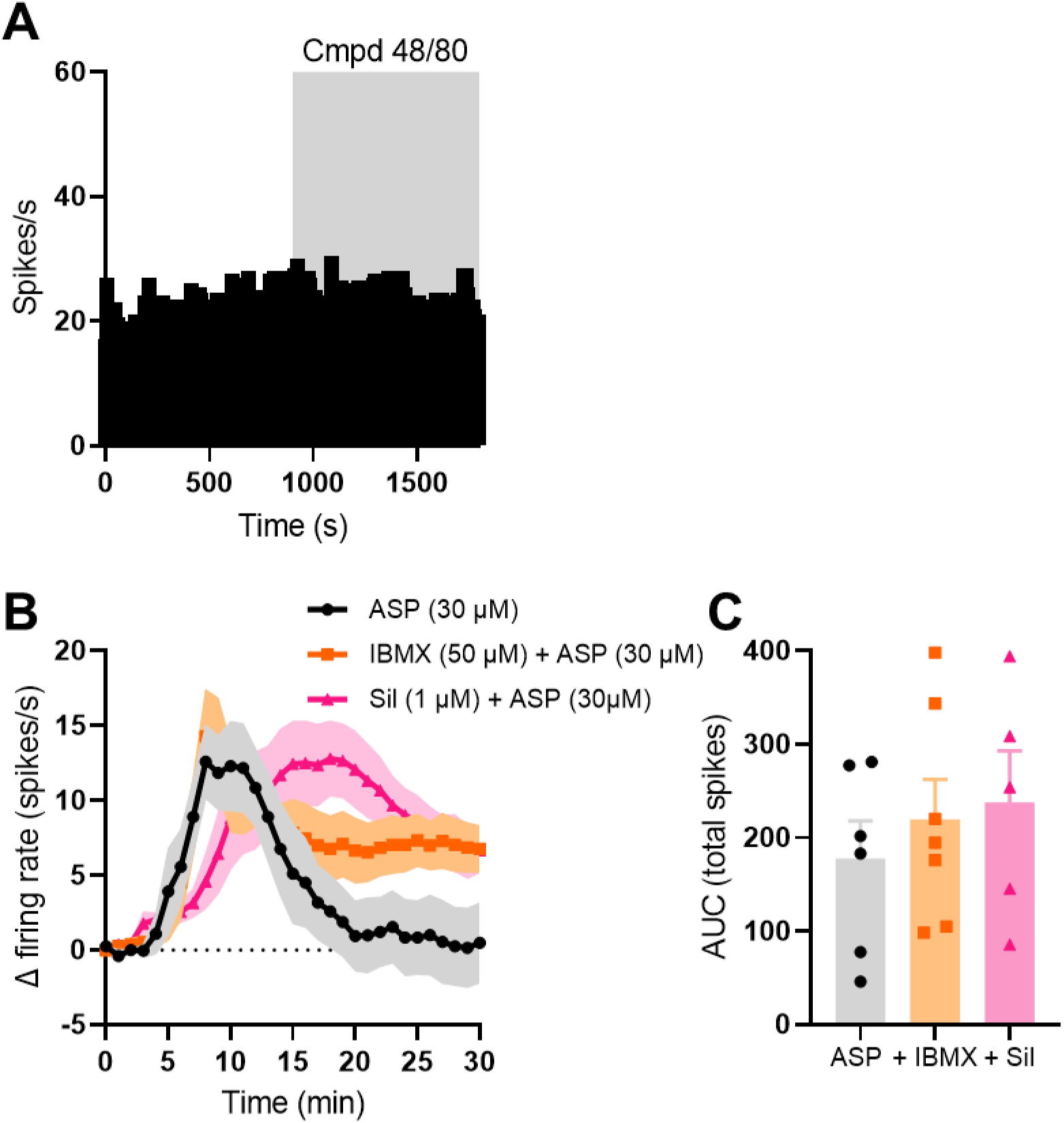
Off-target effects of CS and Zap are unlikely to mediate their inhibition of ASP7663-evoked afferent firing. (A) Example rate histogram showing the afferent firing rate before and during the application of Compound 48/80. (B) Grouped data showing the mean change in afferent firing following the application of ASP7663 (30 µM) to control tissue (black) and tissue pre-incubated with IBMX (50 µM, orange) or sildenafil (1 µM, pink). (C) Grouped data showing the area under the afferent response curves in (B). One-way ANOVA with Tukey’s post-hoc tests.

To verify that the effect of CS and Zap on ASP7663-evoked afferent discharge was mediated by GPR35, we repeated these experiments in tissue from GPR35^-/-^ animals. ASP7663 stimulated similar afferent firing compared to wildtype animals (10.7±1.0 spikes/s, N = 5). However, in GPR35^-/-^ tissue, neither 100 µM CS (8.4±1.3 spikes/s, N = 5, p = 0.51) nor 100 µM Zap (15.5±3.7 spikes/s, N = 5, p = 0.33) had any effect on ASP7663-induced afferent firing (Figure 2E).

CS and Zap display additional pharmacological activity: CS is a mast cell stabiliser (Bernstein et al., 1972) and Zap inhibits phosphodiesterase (PDE) activity (Lugnier et al., 1986). We sought to understand any possible contribution of these GPR35-independent effects to the action of CS and Zap. Consistent with previous reports in healthy colonic tissue, application of the mast cell degranulator, compound 48/80 (50 µg/mL), had no effect on colonic afferent nerve discharge indicating that mast cells were unlikely to have contributed to ASP7663 mediated nerve activation and, consequently, the inhibitory actions of CS (Supplemental Figure 2A). Neither IBMX (50 µM, non-selective PDE inhibitor) nor sildenafil (1 µM, selective PDE5/6 inhibitor) attenuated ASP7663-induced colonic afferent firing (Supplemental Figure 2B and C), suggesting that the suppression of afferent activity by Zap was independent of PDE inhibition.

### Inhibition of TRPA1 but not GPR35 receptor activation attenuates colonic afferent mechanosensitivity

Having established that GPR35 agonists CS and Zap attenuate ASP7663-induced colonic afferent firing, consistent with the marked co-expression of TRPA1 and GPR35 in colonic neurons, we next sought to investigate the effect of GRP35 receptor activation on the afferent response to colorectal distension. Repeated ramp distension of the colon resulted in comparable peak changes in afferent activity after the first ramp, with the afferent firing during ramps two and three being indistinguishable (Supplemental Figure 3A).

TRPA1 has been shown to be an effector of colonic afferent mechanosensitivity and, in keeping with these reports, we confirmed that pre-treatment with a TRPA1-selective antagonist, AM0902 (1 µM), attenuated the colonic afferent response to colonic distension compared to vehicle-treated tissue (main effect of drug, p = 0.077, Figure 3A and D) at distension pressures >35 mmHg (p < 0.041). There was a marked reduction in the peak afferent response to colorectal distension in tissues pre-treated with AM0902 (N = 6) compared to vehicle (N = 8, p = 0.0004, Figure 3E). Application of AM0902 did not affect colonic compliance (Supplemental Figure 3B).

**Figure 3:**
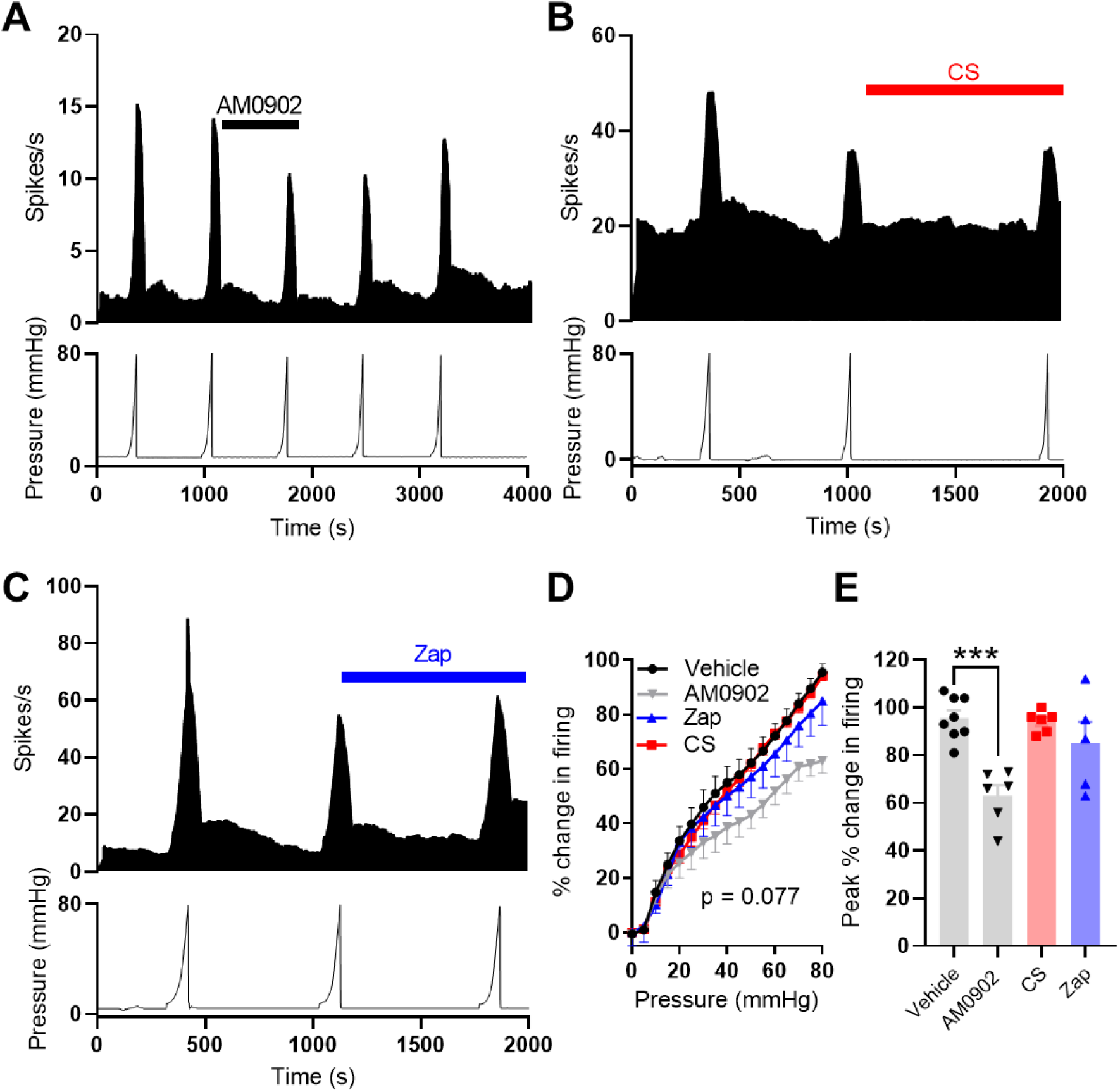
TRPA1 contributes to colonic afferent mechanosensitivity. (A) Example recording showing afferent firing during five successive ramp distensions with AM0902 (1 µM) applied before and during the third ramp distension. (B) Example recording showing afferent firing during three consecutive ramp distensions with CS applied before and during the third ramp distension. (C) Example recording showing afferent firing during three consecutive ramp distensions with Zap applied before and during the third ramp distension. (D) Grouped data showing the percentage change in afferent firing during the third ramp distension in vehicle-(black), AM0902-(grey), CS- (red) and Zap- (blue) treated tissue. Two-way repeated measures ANOVA with Dunnett’s post-hoc tests. The percentage change in firing is relative to peak firing during the preceding ramp distension. (E) Grouped data showing the peak change in afferent firing during the third ramp distension in vehicle-, AM0902-, CS- and Zap-treated tissue. One-way ANOVA with Dunnett’s post-hoc tests.

**Supplemental Figure 3:**
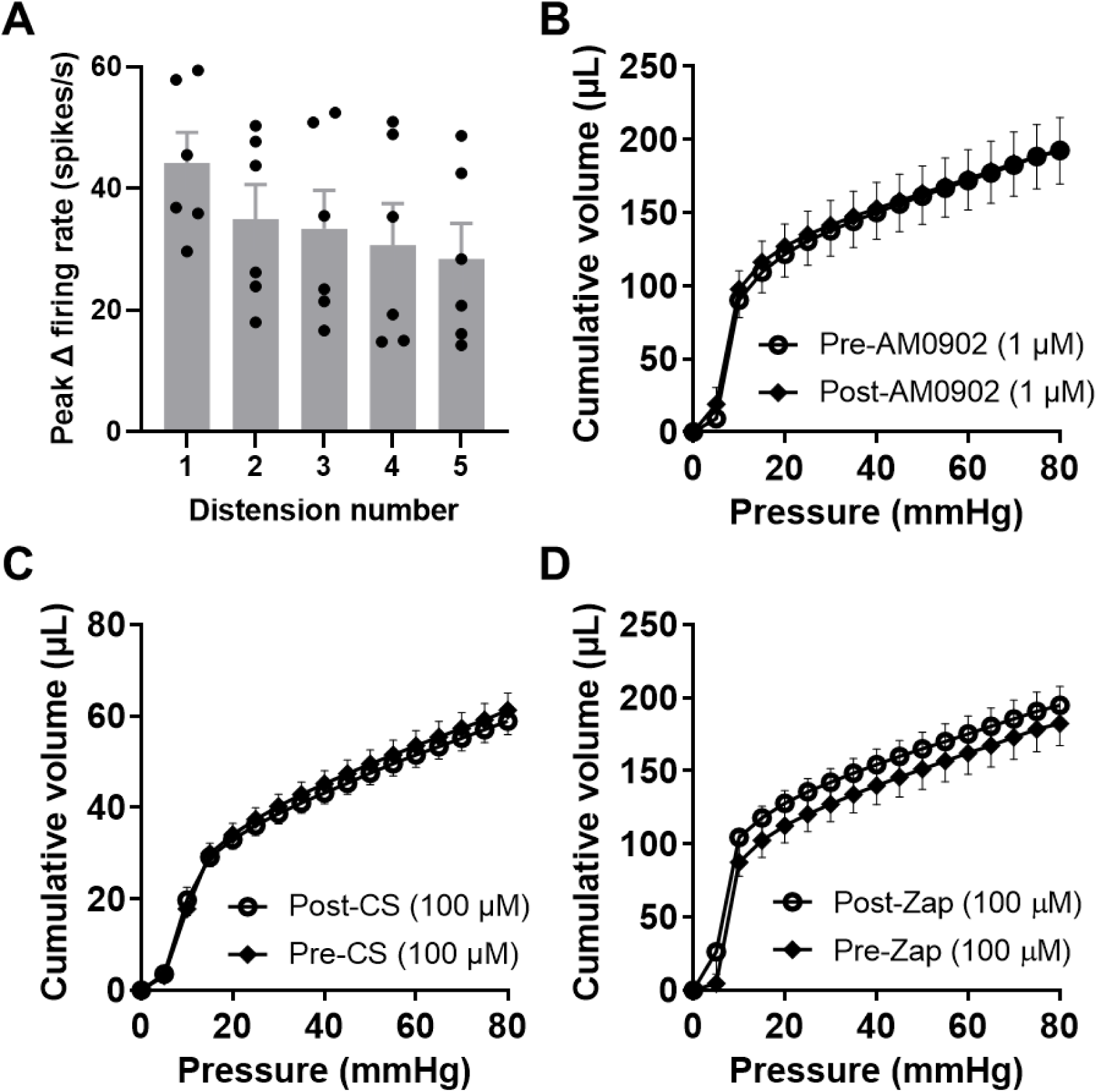
Colonic compliance is not affected by inhibition or activation of TRPA1. (A) Grouped data showing the peak change in afferent firing for each of five consecutive ramp distensions to 80 mmHg. Peak afferent firing during ramps 2 and 3 is indistinguishable (p = 0.87). One-way repeated measures ANOVA (main effect of distension, p = 0.0071; ramp 1 vs ramp 5, p = 0.021; ramp 3 vs ramp 5, p = 0.020). (B) Grouped data showing the volume needed to raise the luminal pressure to 80 mmHg before and after AM0902. Two-way repeated measures ANOVA (main effect of drug, p = 0.90). (C) Grouped data showing the volume needed to raise the luminal pressure to 80 mmHg before and after CS application. Two-way repeated measures ANOVA (main effect of drug, p = 0.69). (D) Grouped data showing the volume needed to raise the luminal pressure to 80 mmHg before and after Zap application. Two-way repeated measures ANOVA (main effect of drug, p = 0.69).

Pre-treatment with CS (100 µM, N = 6) did not affect the colonic afferent response to distension (main effect of drug, p = 0.077, Figure 3B and D) at any intraluminal pressure (p > 0.81). There was no difference in the peak afferent response to colonic distension between vehicle- and CS-treated tissues (p = 0.99, Figure 3E). Zap (100 µM, N = 5) had no effect on the colonic afferent response to ramp distension (main effect of drug, p = 0.077, Figure 3C and D) at any intraluminal pressure (p > 0.32). The peak afferent response to colonic distension was unaffected by Zap (p = 0.31, Figure 3E). Neither the application of CS (Supplemental Figure 3C), nor the application of Zap (Supplemental Figure 3D) had any effect on colonic compliance.

### Activation of GPR35 inhibits ASP7663-induced mechanical hypersensitivity

To explore the relationship between mechanosensitivity, TRPA1 and GPR35 receptor activation further, we investigated the effect of GPR35 agonist pre-treatment on TRPA1 agonist-induced mechanical hypersensitivity. As previously reported, application of ASP7663 (30 µM, Figure 4Ai) augmented the afferent response to colonic distension (main effect of drug, p = 0.016; Figure 4Ai and ii) resulting in an elevated peak increase in afferent firing during ramp distension (10.1±2.3 spikes/s pre-ASP7663 vs 13.2±2.4 spikes/s post-ASP7663, N = 5, p = 0.0004, Figure 4Aiii). Furthermore, the application of ASP7663 elicited a comparable mechanical hypersensitivity in tissue from GPR35^-/-^ mice (main effect of drug, p = 0.0024 Figure 4Bi and ii), resulting in an increase in peak afferent firing during ramp distension from 18.3±2.3 spikes/s to 22.8±2.4 spikes/s (N = 5, p = 0.012, Figure 4Biii). The proportional change in peak distension-evoked afferent firing by ASP7663 was no different between wildtype and GPR35^-/-^ tissue (p = 0.40). No change in colonic compliance was observed following the application of ASP7663 in tissue from wildtype or GPR35^-/-^ mice (Supplemental Figure 4A).

**Figure 4:**
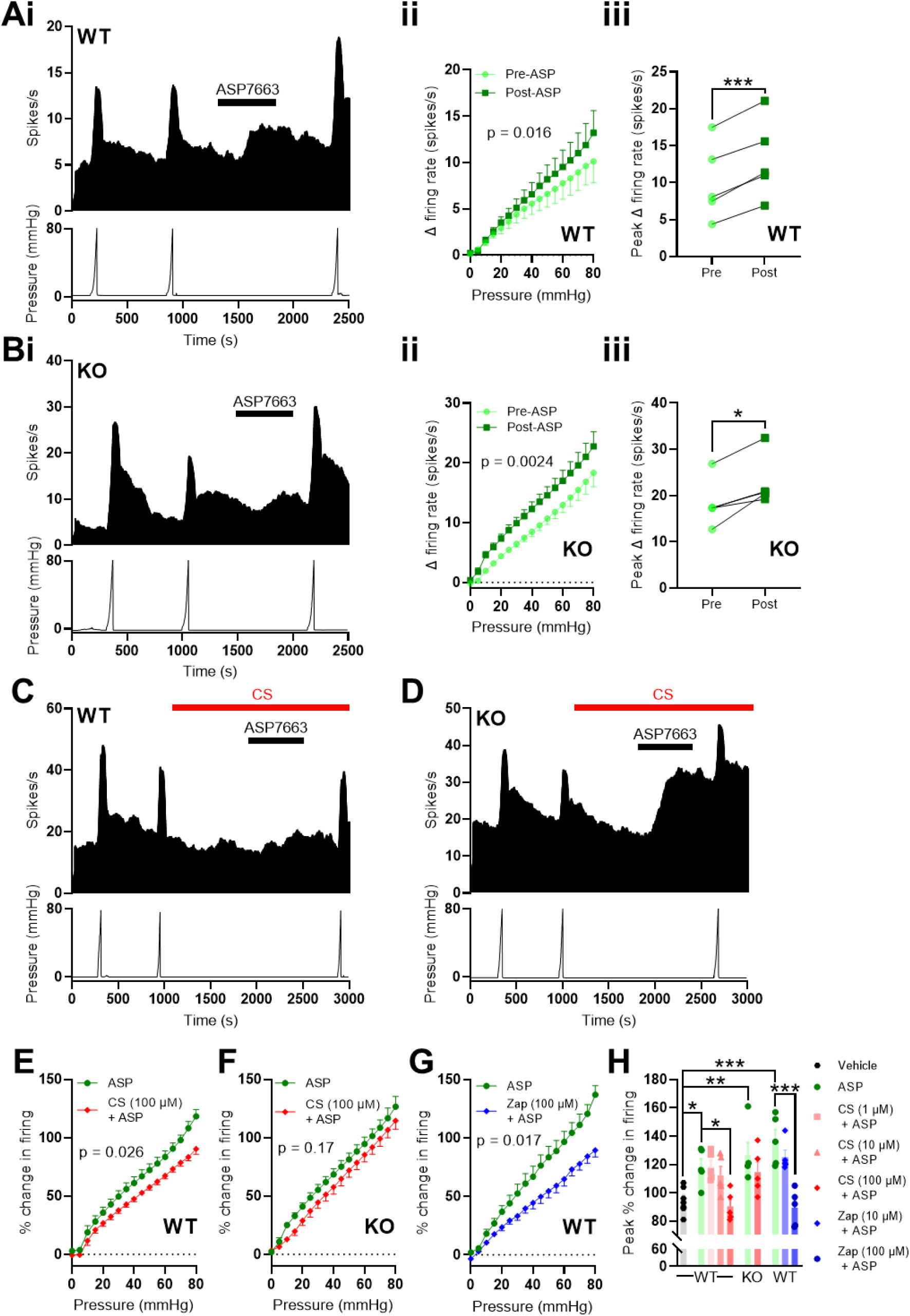
GPR35 activation inhibits TRPA1-induced mechanical hypersensitivity. (A) *(i)* Example recording showing the sensitisation of the afferent response to ramp distension of the colon by ASP7663 in tissue from a wildtype mouse. *(ii)* Grouped data showing the change in afferent firing during ramp distension of the colon before and after ASP7663 application. Two-way repeated measures ANOVA. *(iii)* Grouped data showing the peak change in afferent firing during ramp distension of the colon before and after ASP7663 application. Two-tailed ratio-paired t-test. (B) *(i)* Example recording showing the sensitisation of the afferent response to ramp distension of the colon by ASP7663 in tissue from a GPR35^-/-^ mouse. *(ii)* Grouped data showing the change in afferent firing during ramp distension of the colon before and after ASP7663 application. Two-way repeated measures ANOVA. *(iii)* Grouped data showing the peak change in afferent firing during ramp distension of the colon before and after ASP7663 application. Two-tailed ratio-paired t-test. (C)Example recording from tissue from a wildtype mouse showing the inhibition of ASP7663-induced afferent firing and sensitisation of the response to ramp distension by pre-treatment with CS. (D) Example recording from tissue from a GPR35^-/-^ mouse showing the loss of the inhibitory effect of CS on ASP7663-induced afferent firing and sensitisation of the response to ramp distension. (E) Grouped data showing the change in afferent firing during the third (post-ASP7663) ramp distension in tissue from wildtype mice treated with ASP7663 with or without CS. Two-way repeated measures ANOVA with Holm-Sidak post-hoc tests. (F) Grouped data showing the change in afferent firing during the third (post-ASP7663) ramp distension in tissue from GPR35^-/-^ mice treated with ASP7663 with or without CS. Two-way repeated measures ANOVA with Holm-Sidak post-hoc tests. (G) Grouped data showing the change in afferent firing during the third (post-ASP7663) ramp distension in tissue from wildtype mice treated with ASP7663 with or without Zap. Two-way repeated measures ANOVA with Holm-Sidak post-hoc tests. (H) Grouped data showing the peak change in afferent firing during the third (post-vehicle or -ASP7663) ramp distension. One-way ANOVA with Bonferroni’s post-hoc tests.

**Supplemental Figure 4:**
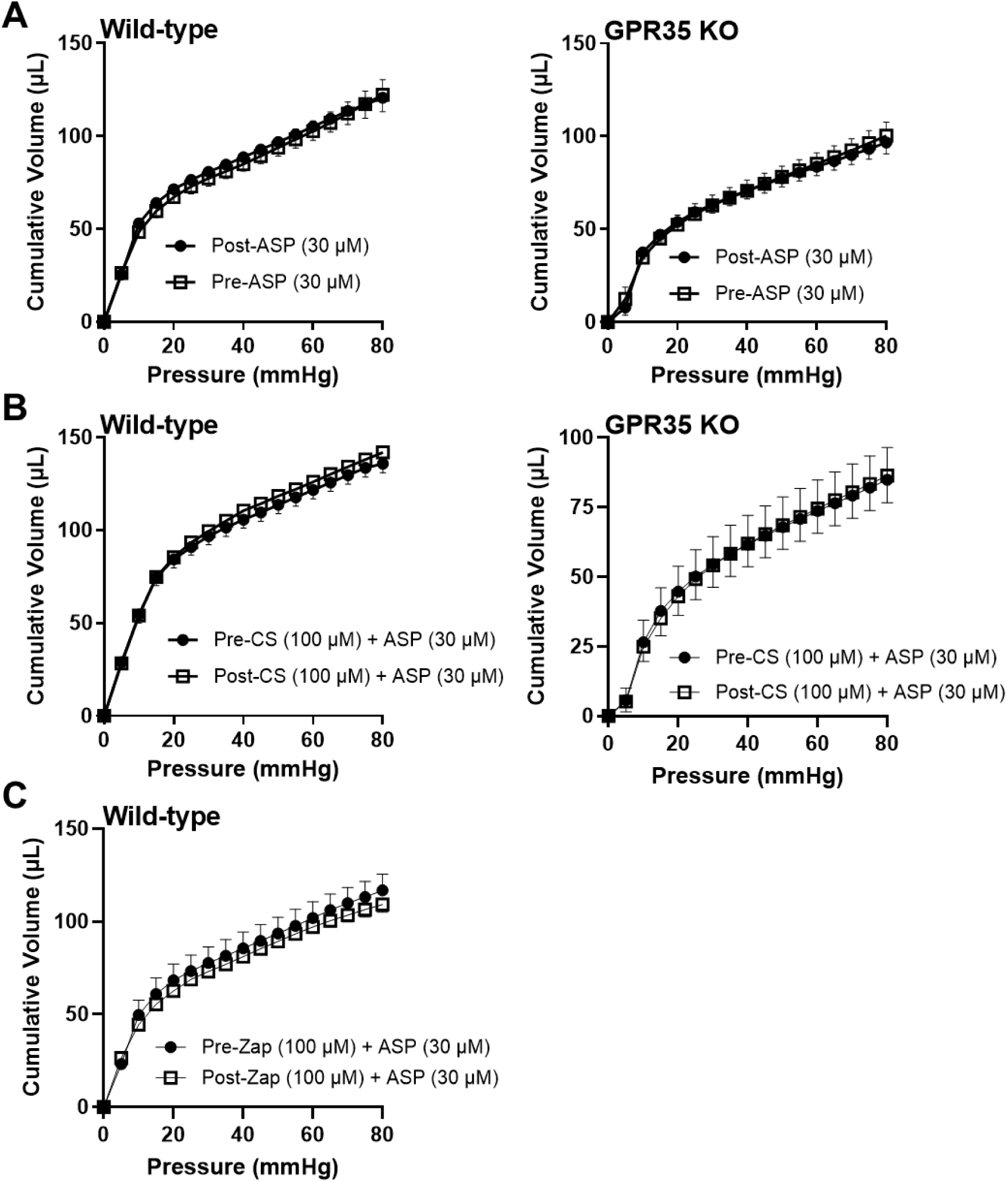
Stimulation of GPR35 does not change the compliance of the colon. (A) *(Left)* Grouped data showing the volume needed to raise the luminal pressure to 80 mmHg before and after ASP7663 application to wildtype tissue. Two-way repeated measures ANOVA (main effect of drug, p = 0.78). *(Right)* Grouped data showing the volume needed to raise the luminal pressure to 80 mmHg before and after ASP7663 application to GPR35^-/-^ tissue. Two-way repeated measures ANOVA (main effect of drug, p = 0.92). (B) *(Left)* Grouped data showing the volume needed to raise the luminal pressure to 80 mmHg before and after CS and ASP7663 application to wildtype tissue. Two-way repeated measures ANOVA (main effect of drug, p = 0.52). *(Right)* Grouped data showing the volume needed to raise the luminal pressure to 80 mmHg before and after CS and ASP7663 application to GPR35^-/-^ tissue. Two-way repeated measures ANOVA (main effect of drug, p > 0.99). (C) Grouped data showing the volume needed to raise the luminal pressure to 80 mmHg before and after Zap and ASP7663 application to wildtype tissue. Two-way repeated measures ANOVA (main effect of drug, p = 0.59).

Consistent with GPR35-mediated inhibition of ASP7663-evoked afferent firing, pre-treatment with CS attenuated ASP7663-induced mechanical hypersensitivity in wildtype (Figure 4C), but not GPR35^-/-^ (Figure 4D), tissue. In wildtype tissue, pre-incubation with 100 µM CS attenuated ASP7663-induced mechanical hypersensitivity (main effect of drug, p = 0.026, Figure 4E). However, 100 µM CS failed to suppress ASP7663-induced mechanical hypersensitivity in tissue from GPR35^-/-^ animals (main effect of drug, p = 0.17, Figure 4F). 100 µM Zap exhibited a similar effect to that of CS and was found to inhibit ASP7663-induced mechanical hypersensitivity in wildtype tissue (main effect of drug, p = 0.017, Figure 4G).

The effect of CS and Zap on the peak change in firing evoked by colonic distension was concentration-dependent. Neither 1 µM (p > 0.99) nor 10 µM (p > 0.99) CS inhibited the effect of ASP7663 on peak distension-evoked afferent activity (Figure 4H), whereas 100 µM CS did suppress the peak change in afferent activity (p = 0.016, Figure 4H). In tissue from GPR35^-/-^ animals, 100 µM CS did not reduce the peak change in distension-evoked afferent firing following ASP7663 application (p > 0.99, Figure 4H). Finally, while 10 µM Zap was without effect (p > 0.99), 100 µM Zap suppressed peak distension-evoked afferent activity following ASP7663 application (p < 0.0001, Figure 4H). Peak distension-evoked firing in each group treated with ASP7663 alone was greater than that in a vehicle-treated group, verifying ASP7663-induced mechanical hypersensitivity (Figure 4H). The application of CS (Supplemental Figure 4B) or Zap (Supplemental Figure 4C) with ASP7663 did not have any effect on colonic compliance.

### Substance P contributes to TRPA1-mediated colonic afferent activation by stimulating sensory neurons

We have thus shown that the GPR35 agonists CS and Zap attenuate TRPA1-induced afferent activation and mechanical hypersensitivity, but not the activation of colonic afferents by luminal distension alone. Therefore, we reasoned that GPR35 receptor activation might block release of neuropeptides from colonic afferent terminals in response to TRPA1 activation rather than having a direct inhibitory effect on the afferent terminal or TRPA1 channel activity, both of which would be expected to attenuate distension-evoked afferent responses. Agonist activation of TRPA1 has previously been shown to produce neurogenic inflammation through the release of SP from sensory neuron terminals, an effect blocked by pre-treatment with agonists of G_i/o_-coupled receptors (Kondo et al., 2005; Engel et al., 2011; Meseguer et al., 2014).

To investigate this, we first confirmed that the gene encoding SP, *Tac1*, is highly co-expressed with *Trpa1* in colonic sensory neurons (Figure 5A). Following this, we demonstrated that pre-treatment with the NK_1_R antagonist, aprepitant (10 µM), blunted the afferent response to ASP7663 (30 µM, Figure 5B), reducing the peak firing elicited by ASP7663 from 12.1±2.9 spikes/s (N = 5) to 4.4±1.6 spikes/s (N = 5, p = 0.048, Figure 5C), indicating that SP and NK_1_R signalling contribute to the afferent response to TRPA1 activation.

**Figure 5:**
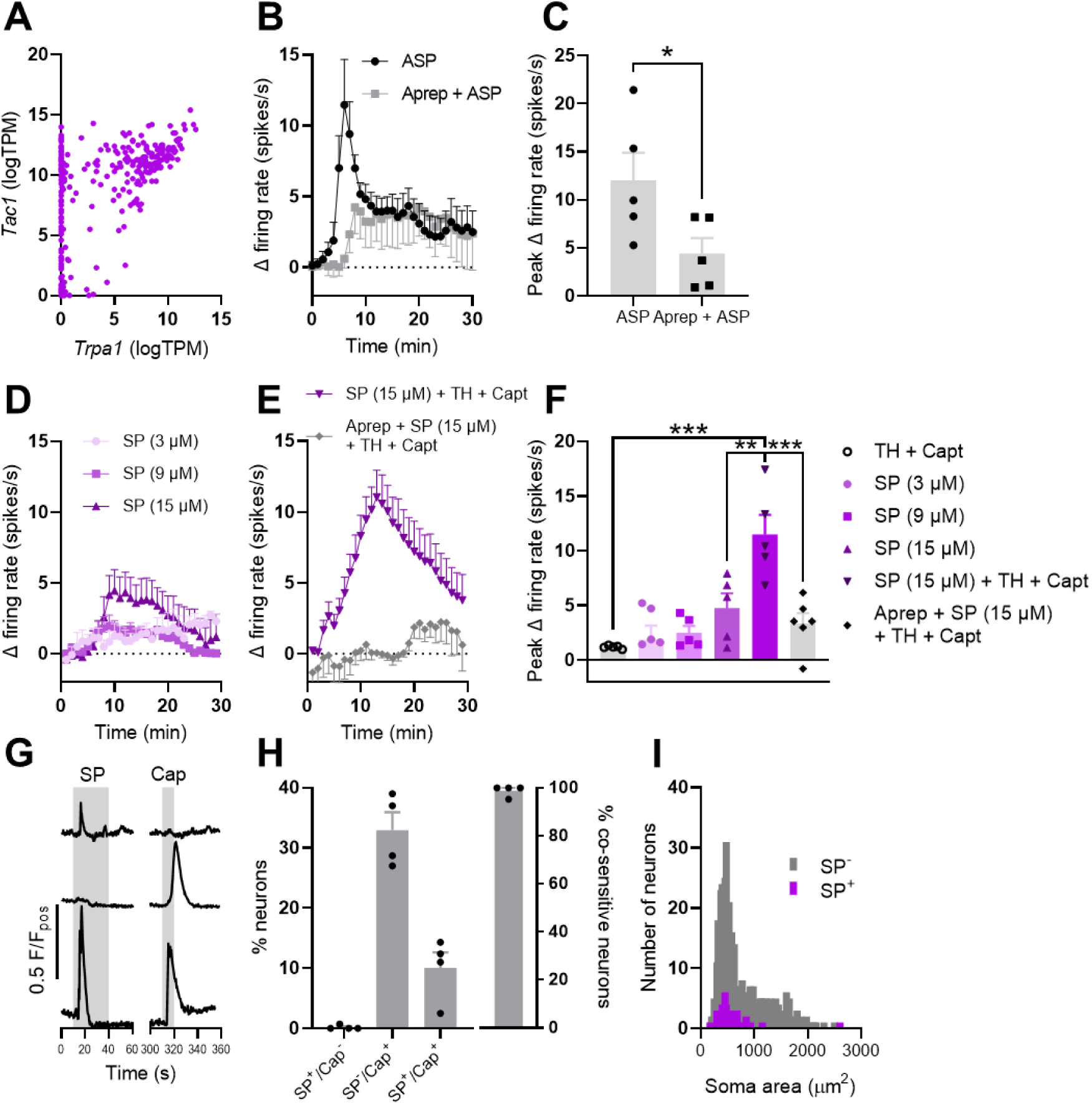
SP stimulates colonic afferents and cultured sensory neurons. (A) Co-expression of *Trpa1* and *Tac1* transcripts in colonic sensory neurons. Data redrawn from Hockley *et al*., 2019. (B) Grouped data showing the change in afferent firing rate following the application of ASP7663 (100 µM) alone or after tissue pretreatment with aprepitant (10 µM). (C) Grouped data showing the peak change in afferent firing rate following the application of ASP7663 (100 µM) alone or after tissue pretreatment with aprepitant (10 µM). Two-tailed unpaired t-test. (D) Grouped data showing the change in afferent firing rate following the application of 10, 30 or 50 µM SP. (E) Grouped data showing the change in afferent firing rate following the application of SP with thiorphan (TH) and captopril (Capt) in the absence and presence of aprepitant. (F) Grouped data showing the peak change in afferent firing rate from the experiments shown in (D) and (E). One-way ANOVA with Bonferroni’s post-hoc tests. (G) Example Fluo-4 fluorescence traces showing three distinct response profiles following the application of SP and capsaicin. (H) *(Left)* Grouped data showing the proportion of neurons which responded to SP alone, capsaicin alone or to both SP and capsaicin. *(Right)* Grouped data showing the proportion of SP-sensitive neurons which were co-sensitive to capsaicin. (I) Histogram showing the distribution of sensory neuron soma size for SP-sensitive (purple) and SP-insensitive (grey) neurons. Median soma areas compared using Mann-Whitney U-test.

**Supplemental Figure 5:**
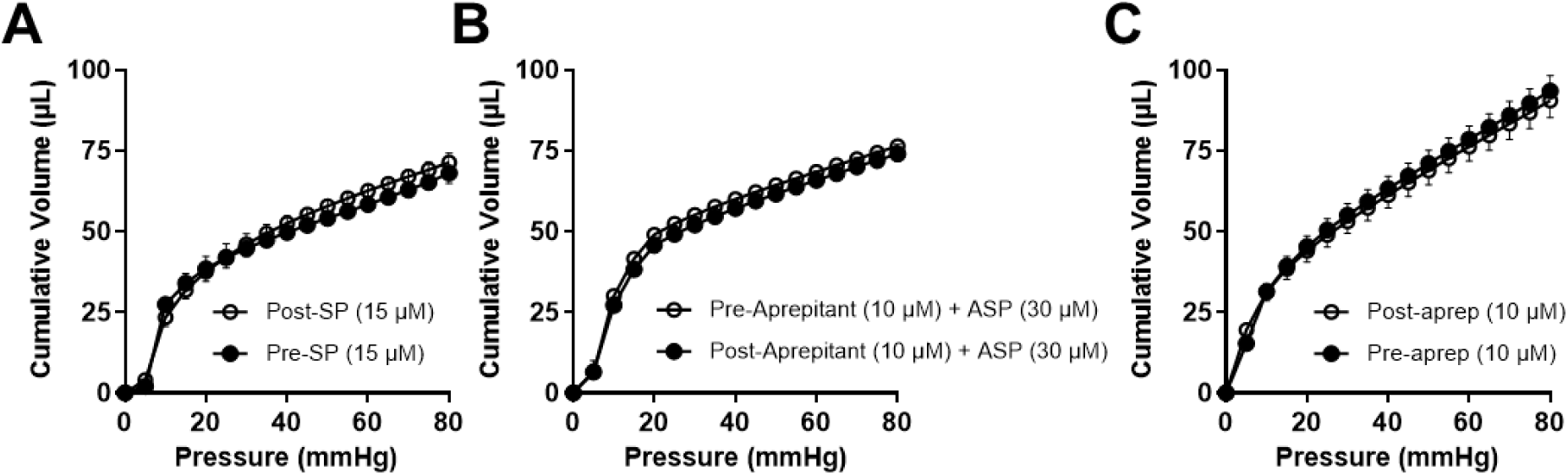
_NK1R_ inhibition does not affect colonic compliance. (A) Grouped data showing the volume needed to raise the luminal pressure to 80 mmHg before and after SP application to wildtype tissue. Two-way repeated measures ANOVA (main effect of drug, p = 0.75). (B) Grouped data showing the volume needed to raise the luminal pressure to 80 mmHg before and after aprepitant and ASP7663 application to wildtype tissue. Two-way repeated measures ANOVA (main effect of drug, p = 0.61). (C) Grouped data showing the volume needed to raise the luminal pressure to 80 mmHg before and after aprepitant application to wildtype tissue. Two-way repeated measures ANOVA (main effect of drug, p = 0.77).

Modest changes in colonic afferent activity were initially found in response to the application of 3 µM, 9 µM and 15 µM SP (Figure 5D), resulting in a peak increase of 1.9±1.3 spikes/s (N = 5), 2.5±0.6 spikes/s (N = 5) and 4.8±1.3 spikes/s (N = 5), respectively (Figure 5F). However, robust increases in colonic afferent activity in response to SP were seen following pre-treatment of tissues with protease inhibitors, thiorphan (10 µM) and captopril (10 µM), highlighting the impact of proteolytic degradation on exogenously applied SP (Figure 5E). The application of 15 µM SP elicited a peak increase in afferent firing of 11.5±1.8 spikes/s (N = 5, p = 0.0035 compared to 15 µM SP alone, Figure 5E and F). The addition of aprepitant abrogated the peak afferent response to SP (3.4±1.0 spikes/s, N = 5, p = 0.0002, Figure 5E and F). The application of thiorphan and captopril alone did not stimulate any appreciable afferent firing (1.2±0.1 spikes/s, N = 5, Figure 5F).

Finally, we further confirmed the stimulatory effect of SP on sensory neurons in culture (where proteases are not likely present), observing a rise in cytosolic [Ca^2+^] in 52 of 497 neurons (from 4 animals) following the application of 100 nM SP. The majority (51 of 52) of SP-sensitive neurons were co-sensitive to the TRPV1 agonist, capsaicin (Figures 5G and H), a key feature of a subset of nociceptive sensory neurons. SP-sensitive neurons were also found to be of a smaller soma size (median soma area: 491 µm^2^; interquartile range: 408-646 µm^2^) compared to SP-insensitive neurons (553 µm^2^, interquartile range: 423-956 µm^2^, p = 0.022, Figure 5I).

### ASP7663-induced mechanical hypersensitivity is dependent on SP signalling

Building on the observation that SP mediates – at least in part – the activation of colonic afferents in response to the stimulation of TRPA1, we next sought to understand the contribution of SP to TRPA1 agonist-induced mechanical hypersensitivity. First, we demonstrated that application of 15 µM SP, in the presence the protease inhibitors thiorphan and captopril, evoked mechanical hypersensitivity comparable to that observed following ASP7663 application (Figure 6A). SP augmented distension-evoked afferent firing (main effect of drug, p = 0.0032, Figure 6B). The peak change in distention-evoked afferent firing was 102.7±4.7% in control conditions (N = 6) compared to 130.8±3.1% following SP application (N = 5, p = 0.0009, Figure 6C).

**Figure 6:**
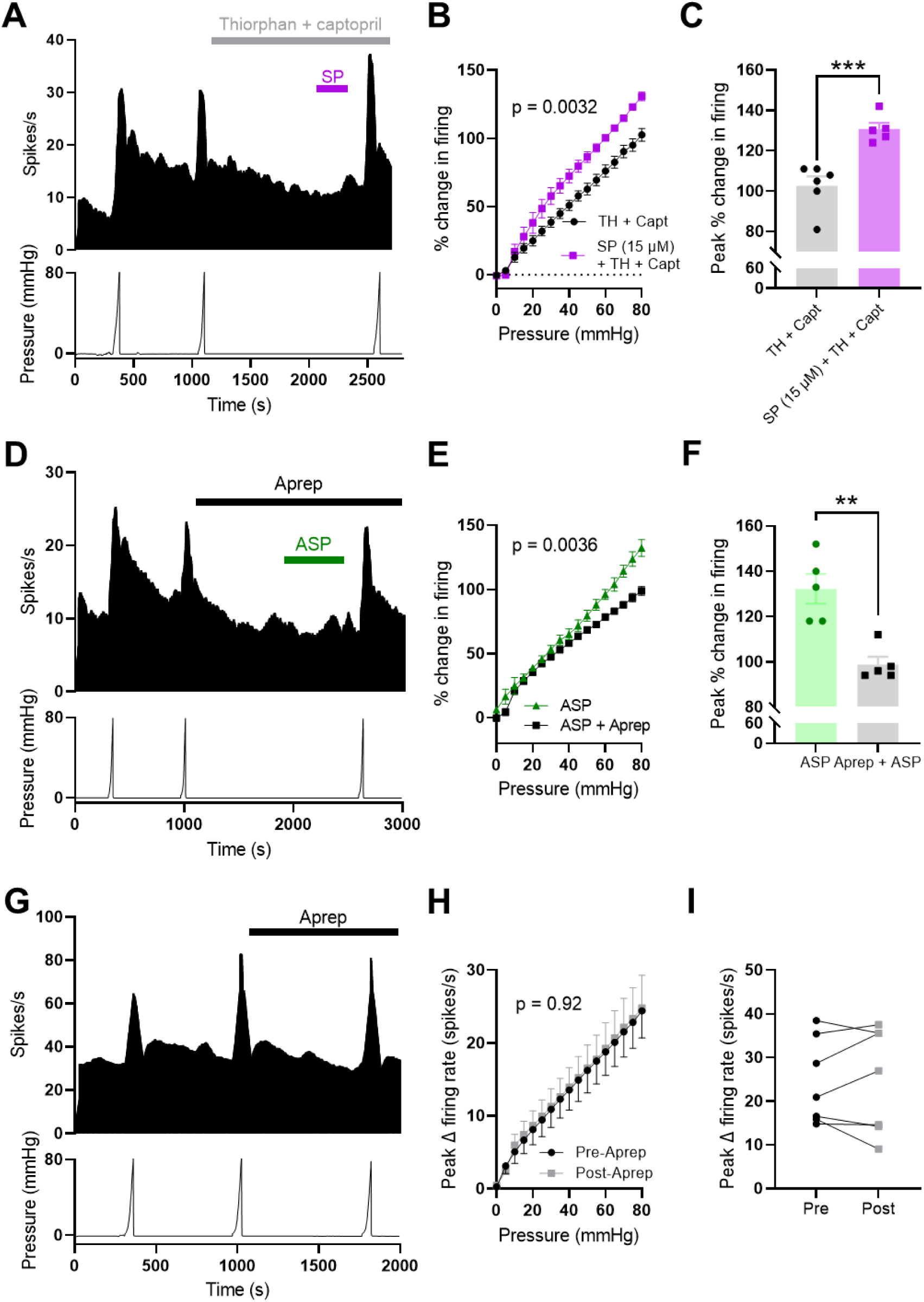
Substance P signalling is required for ASP7663-induced mechanical hypersensitivity. (A) Example recording showing afferent firing during three consecutive ramp distensions of the colon, with SP, thiorphan and captopril applied between the second and third ramps. (B) Grouped data showing the percentage change in afferent firing during the third ramp distension following pretreatment with either thiorphan and captopril alone or thiorphan and captopril with 15 µM SP. Two-way repeated measures ANOVA with Holm-Sidak post-hoc tests. (C) Grouped data showing the peak percentage change in afferent firing from the experiments shown in (A) and (B). Two-tailed unpaired t-test. (D) Example recording showing afferent firing during three consecutive ramp distensions of the colon with aprepitant and ASP7663 applied between the second and third ramps. (E) Grouped data showing the percentage change in afferent firing during the third ramp distension following pretreatment with ASP7663 with or without aprepitant. Two-way repeated measures ANOVA with Holm-Sidak post-hoc tests. (F) Grouped data showing the peak percentage change in afferent firing from the experiments shown in (D) and (E). Two-tailed unpaired t-test. (G) Example recording showing afferent firing during three ramp distensions with aprepitant alone applied between the second and third ramps. (H) Grouped data showing the change in afferent firing rate during the second (pre-aprepitant) and third (post-aprepitant) ramp distensions. Two-way repeated measures ANOVA with Holm-Sidak post-hoc tests. (I) Grouped data showing the peak change in afferent firing rate during the second (pre-aprepitant) and third (post-aprepitant) ramp distensions. Two-tailed paired t-test.

Furthermore, pre-treatment with 10 µM aprepitant ameliorated ASP7663-evoked mechanical hypersensitivity (main effect of drug, p = 0.0036, Figure 6D and E), reducing the peak change in distension-evoked afferent firing from 132.2±6.5% (N = 5) to 94.0±3.4% (N = 5, p = 0.0019, Figure 6F). Treatment with aprepitant alone had no effect on distension-evoked firing (main effect of drug, p = 0.92, Figure 6G and H). There was no difference in peak distension-evoked afferent firing before and after the application of aprepitant (N = 7, p = 0.84, Figure 6I). Treatment with SP, aprepitant and ASP7663, or aprepitant alone did not affect colonic compliance (Supplemental Figure 5).

### Cromolyn blocks TRPA1-mediated SP release and colonic contractility via GPR35

Finally, having identified a role for SP in TRPA1-mediated activation of colonic afferents and mechanical hypersensitivity, we sought to confirm that TRPA1 activation evokes SP release in colonic tissue and whether this could be attenuated by agonist stimulation of GPR35. To do so, we first directly measured SP release from the colon in response to ASP7663 using a chemiluminescent immunoassay (Figure 7A). Treatment of wildtype tissue with ASP7663 (30 µM) elicited a ∼10-fold increase in detected SP compared to the blank control (N = 10 per group, p < 0.0001, Figure 7B). This was blocked by AM0902 (3 µM, N = 8, p = 0.0002, Figure 7B), confirming it is mediated by TRPA1 activation. ASP7663-evoked SP release was also abolished by pre-treatment with CS (100 µM, N = 6, p < 0.0001, Figure 7B).

**Figure 7:**
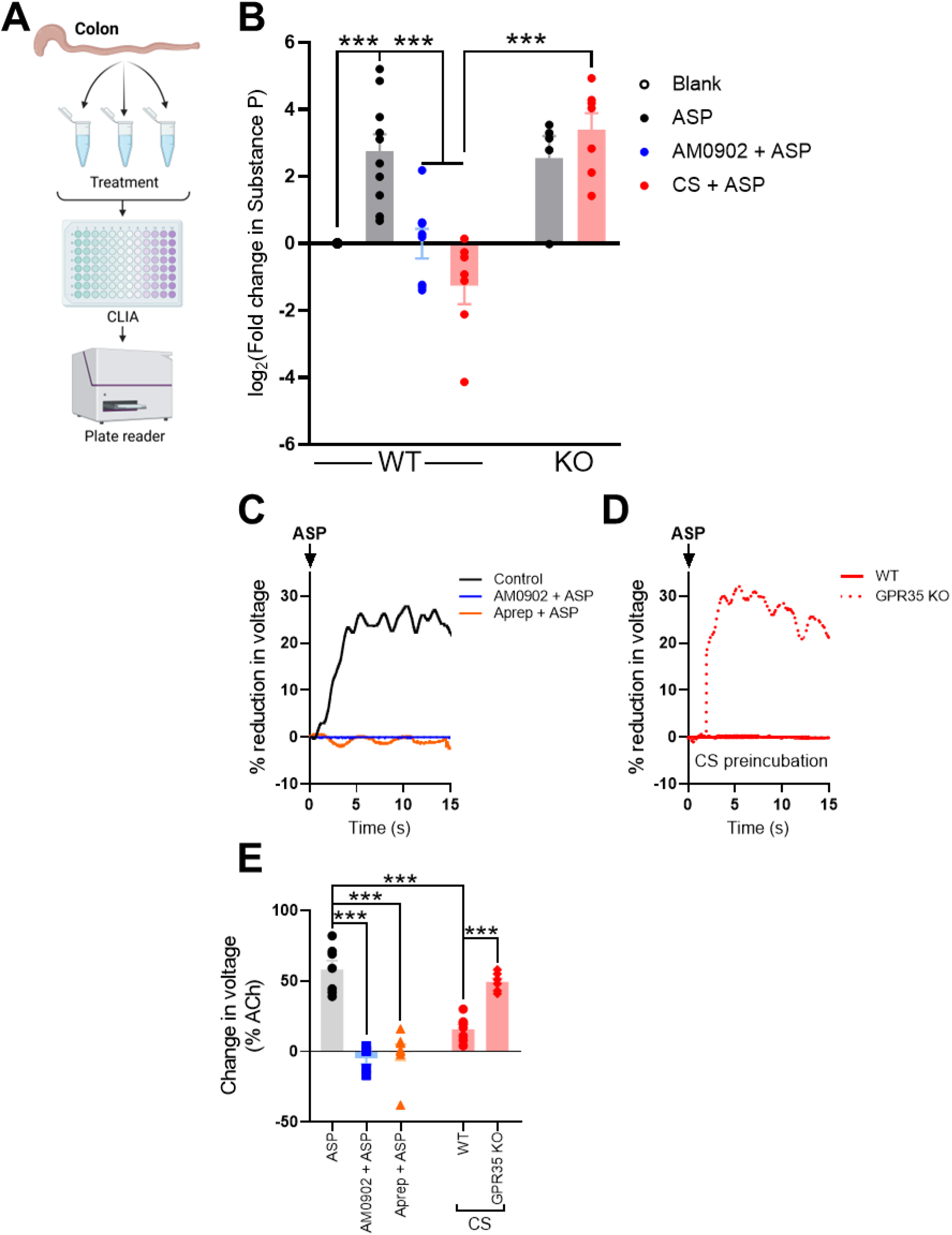
TRPA1 activation evokes SP release in the colon. (A) Schematic showing the experimental protocol for measuring SP release in the colon. (B) Grouped data showing the fold change in SP (relative to a black control experiment) following incubation of tissue with ASP7663 alone or with AM0902 or CS in wildtype and GPR35^-/-^ tissue. One-way ANOVA with Bonferroni’s post-hoc tests. (C) Example recordings showing the reduction in colon length by ASP7663 (shown as a reduction in voltage across a transducer attached to one end of the colon; voltages are shown relative to that evoked by 10 µM ACh). The effect of ASP7663 was blocked by both AM0902 and aprepitant. (D) Example recordings showing the effect of CS pretreatment on ASP7663-induced colonic contraction in tissue from wildtype and GPR35^-/-^ animals. CS failed to inhibit ASP7663-induced contraction in tissue lacking GPR35. (E) Grouped data showing the peak percentage change in transducer voltage (relative to ACh) for the experiments shown in (C) and (D). One-way ANOVA with Bonferroni’s post-hoc tests.

By contrast, CS had no effect on the magnitude of SP release evoked by ASP7663 in tissue from GPR35^-/-^ mice. ASP7663-induced SP release was comparable in the presence (N = 7) or absence (N = 5) of CS (p > 0.99, Figure 7B). ASP7663 induced a similar magnitude of SP release in tissue from wildtype and GPR35^-/-^ mice (p > 0.99, Figure 7B).

Consistent with the findings from the SP release assay, the application of ASP7663 evoked a marked contraction of the mouse distal colon (N = 7, Figure 7C) which was abolished by pre-treatment with either AM0902 (3 µM, N = 5, p < 0.0001, Figure 7C and E) and aprepitant (10 µM, N = 8, p < 0.0001, Figure 7C and E), consistent with the response being mediated by TRPA1-evoked SP release. Pre-treatment with 100 µM CS robustly inhibited ASP7663-evoked colonic contraction in tissue from wildtype mice (N = 7, p < 0.0001 compared to ASP7663 alone, Figure 7D and E). CS had no effect on the contractile response to ASP7663 application in tissue from GPR35^-/-^ mice (N = 6, p > 0.99 compared to ASP7663 alone in wildtype tissue, Figure 7D and E). The effect of CS on ASP7663-evoked colonic contraction was markedly different in tissue from wildtype and GPR35^-/-^ animals (p = 0.0003, Figure 7E); the inhibitory effect of CS was lost in GPR35^-/-^ tissue.

## Discussion

The development of non-opioid analgesics for the treatment of chronic abdominal pain is a pressing area of unmet clinical need. To address this, we examined the distribution of G_i/o_-coupled receptors in colonic sensory neurons to identify drug targets which have the potential to inhibit nociceptor activation, but lack the abuse liability, sedation and gut-related side effects associated with agonists of opioid, cannabinoid and GABA_B_ receptors (Srinath et al., 2014). This led to the identification of several G_i/o_-coupled GPCRs as putative visceral analgesic drug target due to their marked co-expression with TRPA1, a mediator of noxious mechanosensitivity in the bowel (Brierley et al., 2009). We examined the anti-nociceptive effects of GPR35 due to the commercial availability of GPR35 agonists, such as CS and Zap, and GPR35^-/-^ mice.

Consistent with our *in silico* docking studies which identified binding sites for CS and Zap at the GPR35 receptor, we demonstrated that the anti-nociceptive effects of these drugs on TRPA1-mediated colonic nociceptor activation and mechanosensitisation were dependent upon GPR35. Further work showed these anti-nociceptive effects occurred through the inhibition of SP release and by doing so revealed both the pro-nociceptive effect of SP on colonic afferents and the contribution of SP to TRPA1-mediated colonic nociceptor activation. In addition, our data identify GPR35, and its inhibition of SP-mediated colonic contractility and nociceptor firing, as a putative mechanism of action for the reported clinical efficacy of CS in the treatment of IBS (Stefanini et al., 1992 and 1995; Daryani et al., 2009; Carroll et al., 2013). These findings highlight the utility of GPR35 agonists for the treatment of abdominal pain in IBS, as well as other GI diseases, such as IBD.

The findings from our study are strongly supported by the available literature on transcript expression in sensory neurons in the DRG. Previous reports show marked expression of GPR35 in sensory neurons (Ohshiro et al., 2008; Cosi et al., 2011; Hockley et al., 2019; Marti-Solano et al., 2020; Wangzhou et al., 2020), as well as its co-expression with TRPA1 and SP (Hockley et al., 2019). Importantly, this co-expression is conserved in human DRG in which GPR35 is highly expressed in nociceptor populations, including those with enriched expression of TRPA1 (Tavares-Ferreira et al., 2022). Although not reported in the current study, GPR35 also shows marked co-expression with other channels responsible for nociceptive signalling in colonic afferents, such as TRPV1, Na_V_1.8, tropomyosin receptor kinase A (TrkA), histamine H_1_ receptors and 5-HT_3_ receptors (Ohshiro et al., 2008; Cosi et al., 2011; Franck et al., 2011; Egerod et al., 2018; Kim et al., 2023). As such, it is likely that the inhibitory effect of GPR35 stimulation on TRPA1-mediated colonic afferent activation and mechanical hypersensitivity would be replicated across a broader range of noxious and disease-derived stimuli such as acidity, NGF, histamine and 5-HT.

In the present study, we used TRPA1-induced afferent activity and hypersensitivity to study the anti-nociceptive effects of GPR35 on colonic afferent activation due to its established role in noxious colonic mechanosensation (Yang et al., 2008; Kondo et al., 2009; Brierley et al., 2009; Vermeulen et al., 2013). In addition, TRPA1 is a downstream effector for the development of visceral mechanical hypersensitivity in response to a broad range of algogenic stimuli implicated in IBS and IBD (Campaniello et al., 2017). These include inflammatory mediators such as bradykinin (Bandell et al., 2004; Bautista et al., 2006), histamine (Balemans et al., 2019, prostaglandins (PGE_2_) (Dall’Acqua et al., 2014), interleukins (IL-1β) and TNFα (Hughes et al., 2013; Barker et al., 2022), as well as inflammatory soups/supernatants generated from animal models of visceral hypersensitivity, and IBS and IBD biopsy samples (Yang et al., 2008; Hughes et al., 2013). We confirmed the pro-nociceptive potential of TRPA1 by showing that pre-treatment with a selective TRPA1 antagonist, AM0902, inhibited the colonic afferent response to luminal distension at noxious distending pressures and demonstrated that pre-treatment with the TRPA1 agonist ASP7663 produced mechanical hypersensitivity.

Interestingly, the effect of GPR35 activation showed evidence of stimulus selectivity, inhibiting TRPA1-mediated colonic afferent activation and mechanosensitisation but not the response to luminal distension at noxious pressures known to be dependent on TRPA1. Further study revealed this was due to inhibition of SP released following agonist stimulation of TRPA1, which contributed significantly to ASP7663-induced colonic afferent activation and mechanosensitisation as revealed by the inhibition of these effects by aprepitant. However, aprepitant did not suppress the basal afferent response to colonic distension, indicating that SP release does not contribute to distension-evoked afferent firing. This is most likely a consequence of the duration of afferent and TRPA1 channel activity during colonic distension being insufficient to trigger substantive antidromic SP release from nerve terminals. Sustained agonist-induced activity of TRPA1 is seemingly better suited to evoking SP release (Trevisani et al., 2007; Engel et al., 2011; Meseguer et al., 2014). Building on this observation, it may be expected that GPR35 agonists would have the desirable effect of preventing the prolonged activation of nociceptors associated with visceral hypersensitivity and chronic pain in diseases such as IBS and IBD without blocking acute abdominal pain arising in response to potentially life-threatening obstruction of the bowel. Consistent with this hypothesis, CS has previously been reported to reduce symptom severity in IBS patients (Stefanini et al., 1992 and 1995; Daryani et al., 2009), an effect our findings suggest is mediated by GPR35 receptor activation.

In addition to inhibiting TRPA1-mediated colonic afferent activation and mechanosensitisation, SP release and colonic contraction – consistent with the marked expression of GPR35 in colonic sensory neurons – CS has also been reported to have several desirable actions on immune cell function (Puzzovio et al., 2022). Most notably, CS acts as a mast cell stabiliser which is also likely to inhibit colonic afferent activation and pain in disease states such as IBS and IBD (Bernstein et al., 1972; Stefanini et al., 1992 and 1995; Barbara et al., 2004; Klooker et al., 2010). Our data suggest that mast cells are not responsible for the inhibitory effects of CS seen in the current study as responses were replicated with Zap which is not a mast cell stabiliser (Carroll et al., 2013). Furthermore, experiments were also performed using tissue from healthy mice in which colonic mast cell levels are typically low and, as such, unlikely to contribute significantly to changes in colonic afferent activity. This was confirmed by the inability of compound 48/80 to stimulate colonic afferent firing, consistent with previous findings from other groups (Coldwell et al., 2007). Nevertheless, the ability of CS to suppress colonic afferent activity, colonic contractility, and the release of pro-inflammatory and algogenic mediators from mast cells is likely to result in greater clinical efficacy due to the targeting of multiple aspects of disease pathology.

The findings from our study highlight the potential of GPR35 agonists to provide non-opioid-based analgesia in GI diseases, such as IBS and IBD, due to their ability to prevent neurogenic colonic afferent activation, mechanosensitisation and colonic contractility through the inhibition of SP release. These findings shed new light on the mechanism underpinning the clinical efficacy of CS in the treatment of IBS and support the further development of GPR35 agonists for the treatment of GI disorders.

